# Noise-robust recognition of objects by humans and deep neural networks

**DOI:** 10.1101/2020.08.03.234625

**Authors:** Hojin Jang, Devin McCormack, Frank Tong

## Abstract

Deep neural networks (DNNs) for object classification have been argued to provide the most promising model of the visual system, accompanied by claims that they have attained or even surpassed human-level performance. Here, we evaluated whether DNNs provide a viable model of human vision when tested with challenging noisy images of objects, sometimes presented at the very limits of visibility. We show that popular state-of-the-art DNNs perform in a qualitatively different manner than humans – they are unusually susceptible to spatially uncorrelated white noise and less impaired by spatially correlated noise. We implemented a noise-training procedure to determine whether noise-trained DNNs exhibit more robust responses that better match human behavioral and neural performance. We found that noise-trained DNNs provide a better qualitative match to human performance; moreover, they reliably predict human recognition thresholds on an image-by-image basis. Functional neuroimaging revealed that noise-trained DNNs provide a better correspondence to the pattern-specific neural representations found in both early visual areas and high-level object areas. A layer-specific analysis of the DNNs indicated that noise training led to broad-ranging modifications throughout the network, with greater benefits of noise robustness accruing in progressively higher layers. Our findings demonstrate that noise-trained DNNs provide a viable model to account for human behavioral and neural responses to objects in challenging noisy viewing conditions. Further, they suggest that robustness to noise may be acquired through a process of visual learning.

## Introduction

A central question in cognitive and computational neuroscience concerns how we detect, discriminate and identify stimuli by sight [1–3]. The task of object recognition is exceedingly complex, yet human observers can typically recognize most any object within just fractions of a second [4, 5]. The human visual system processes information in a hierarchically organized manner, progressing from the encoding of basic visual features in early visual areas to the representation of more complex object properties in higher visual areas [6–10]. What are the neural computations performed by the visual system that allow for successful recognition across a diversity of contexts and viewing conditions?

There is growing evidence to indicate that deep neural networks (DNNs) trained on object classification provide the best current model of the human and non-human primate visual systems [11, 12]. The visual representations learned by these DNNs demonstrate a reliable correspondence with the neural representations found at multiple levels of the human visual pathway [13–16]. Moreover, DNNs trained on large datasets of object images, such as ImageNet [17], can reliably predict how individual neurons in the monkey inferotemporal cortex will respond to objects, faces, and even synthetic stimuli [18–21].

Although DNNs can perform remarkably well on tasks of object recognition, with claims that they have achieved or even surpassed human-level performance [22, 23], a conundrum lies in the fact that these networks tend to lack robustness to more challenging viewing conditions. In particular, there is some evidence to suggest that DNNs are unusually susceptible to visual noise and clutter, and that human recognition performance is more robust to noisy viewing conditions [24–27]. Identifying potential disparities between human and DNN performance is necessary to understand the limitations of current DNN models of human vision [28, 29] and a precursor to developing better models.

The goal of our study was to determine whether DNNs can provide a viable model of human behavioral and neural performance under stress-test visual conditions. Our recognition task required humans and DNNs to classify objects embedded in either spatially independent noise or spatially correlated noise, across a wide range of signal-to-noise ratios (**Figure 1a**). Spatially independent Gaussian noise has been used to characterize attentional modulation of visual sensitivity and the perceptual learning of complex stimuli [30, 31]. Pixelated noise has also been used to characterize the robustness of visual cortical responses to objects presented in high levels of noise [32]. We were also interested in assessing the impact of Fourier phase-scrambled noise on object recognition performance, as such noise preserves the 1/F amplitude spectrum of natural images [33] and contains spatially correlated structure that might be more confusing to object recognition systems.

**Figure 1.**
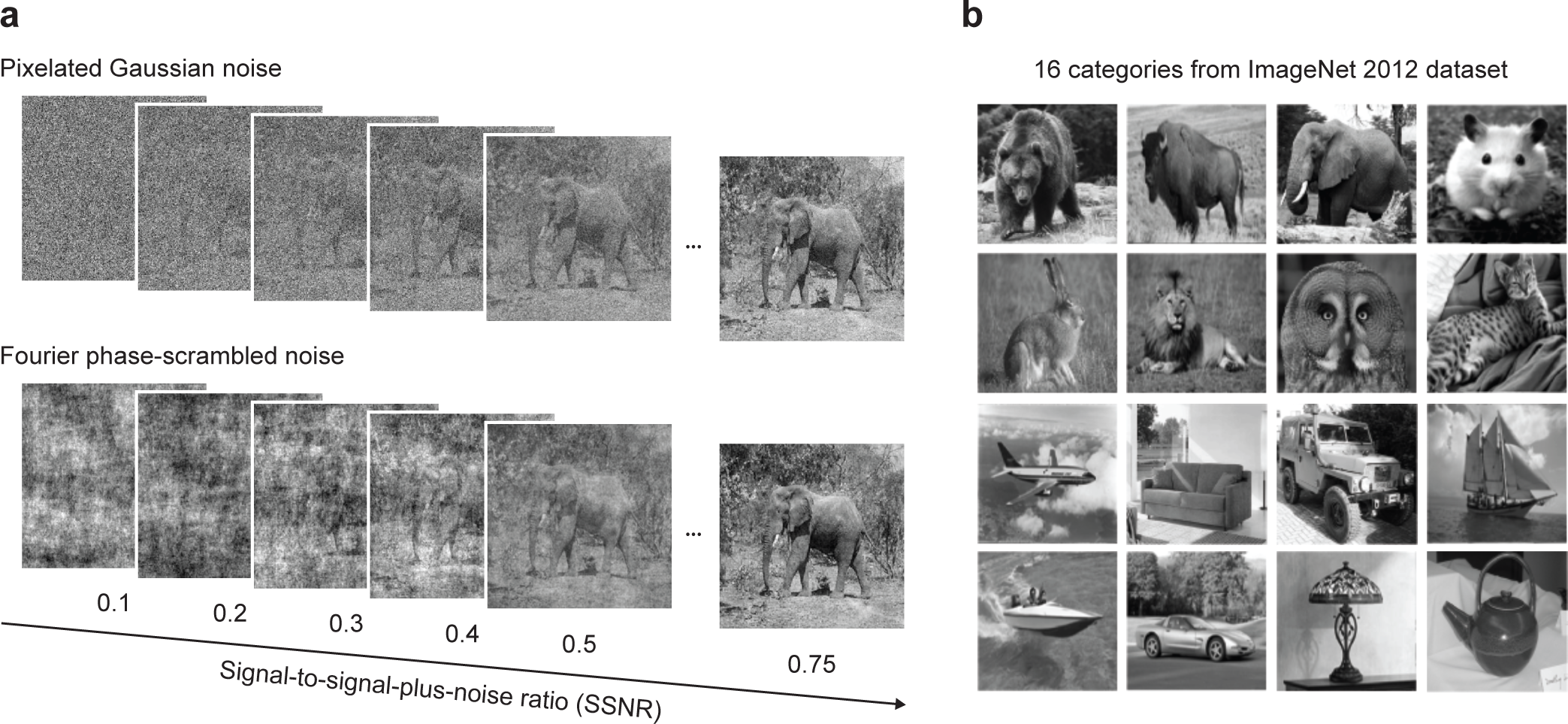
**a** Examples of an object image in pixelated Gaussian noise or Fourier phase-scrambled noise, shown at varying SSNR levels. **b** Example images from the 16 object categories used in this study: bear, bison, elephant, hamster, hare, lion, owl, tabby cat, airliner, couch, jeep, schooner, speedboat, sports car, table lamp, teapot.

Both human and DNN systems can be stressed by lower signal-to-noise ratios, as the object images approach the limits of perceptual visibility. This experimental design allowed us to test for quantitative differences in performance, by identifying the critical noise level at which performance sharply declines. Moreover, it allowed us to test for qualitative differences in visual processing across noise type. We found that popular state-of-the-art DNNs perform in a qualitatively different manner than humans. Specifically, DNNs are unusually susceptible to pixelated Gaussian noise (i.e., white noise) and less susceptible to spatially correlated Fourier phase-scrambled noise (similar to ‘pink’ noise), whereas human observers show the opposite pattern of performance.

Next, we sought to investigate whether DNNs can be trained to recognize objects in extreme levels of visual noise, and in particular, whether such noise-trained DNNs might provide a better match to human behavioral and neural performance. Although it has long been known that neural networks can be regularized by adding a small amount of noise to their input data [34] and such procedures have proven useful in the training of DNNs [35], the impact of training DNNs with extreme levels of noise has only recently begun to receive attention [24–27]. Here, we found that noise-trained DNNs could accurately classify objects at lower noise thresholds than human observers, but more importantly, their qualitative pattern of performance to different noise types provided a better match to human performance. Moreover, noise-trained DNNs performed far better than standard DNNs in their ability to predict human recognition thresholds on an image-by-image basis. A layer-specific analysis of DNN activity patterns indicated that noise training led to widespread changes in the robustness of the network, with more pronounced differences between standard and noise-trained networks found in the middle and higher layers.

We performed a functional neuroimaging experiment to assess the degree of correspondence between DNN models and human neural activity. Multivariate decoding of activity patterns in the human visual cortex revealed better discrimination of objects in pixelated Gaussian noise as compared to Fourier phase-scrambled noise, consistent with the behavioral advantage shown by human observers and also by noise-trained DNNs. Moreover, noise-trained DNNs provided a better correspondence to the patterns of object-specific responses found in both early visual areas and high-level object areas. We go on to show that DNNs trained to recognize objects in artificial noise can generalize their knowledge to some extent to other image distortions, including real-world conditions of visual noise. Taken together, our findings demonstrate that noise-trained DNNs provide a viable model of the noise-robust properties of the human visual system.

## Results

In Experiment 1, we evaluated the performance of 8 pre-trained DNNs (AlexNet, VGG-F, VGG-M, VGG-S, VGG-16, VGG-19, GoogLeNet, and ResNet-152 [36–39]) and 20 human observers at recognizing object images presented in either pixelated Gaussian noise or Fourier phase-scrambled noise (**Figure 1a**, see **Methods**). Object images were presented with varying levels of visual noise by manipulating the signal-to-signal-plus-noise ratio (SSNR), which is bounded between 0 (noise only) and 1 (signal only). This allowed us to quantify changes in performance accuracy as a function of SSNR level. Performance was assessed using images from 16 object categories (**Figure 1b**) obtained from the validation data set of the ImageNet database [17].

These images were novel to the participants and were never used for DNN training.

**Figure 2a** shows the mean performance accuracy of DNNs and humans plotted as a function of SSNR level, with the performance of individual DNNs shown in **Figure 2b**. Although DNNs could match the performance of human observers under noise-free conditions, consistent with previous reports [22], DNN performance became severely impaired in the presence of moderate levels of noise. Most DNNs exhibited a precipitous drop in recognition accuracy as SSNR declined from 0.6 to 0.4, whereas human performance was much more robust across this range.

**Figure 2.**
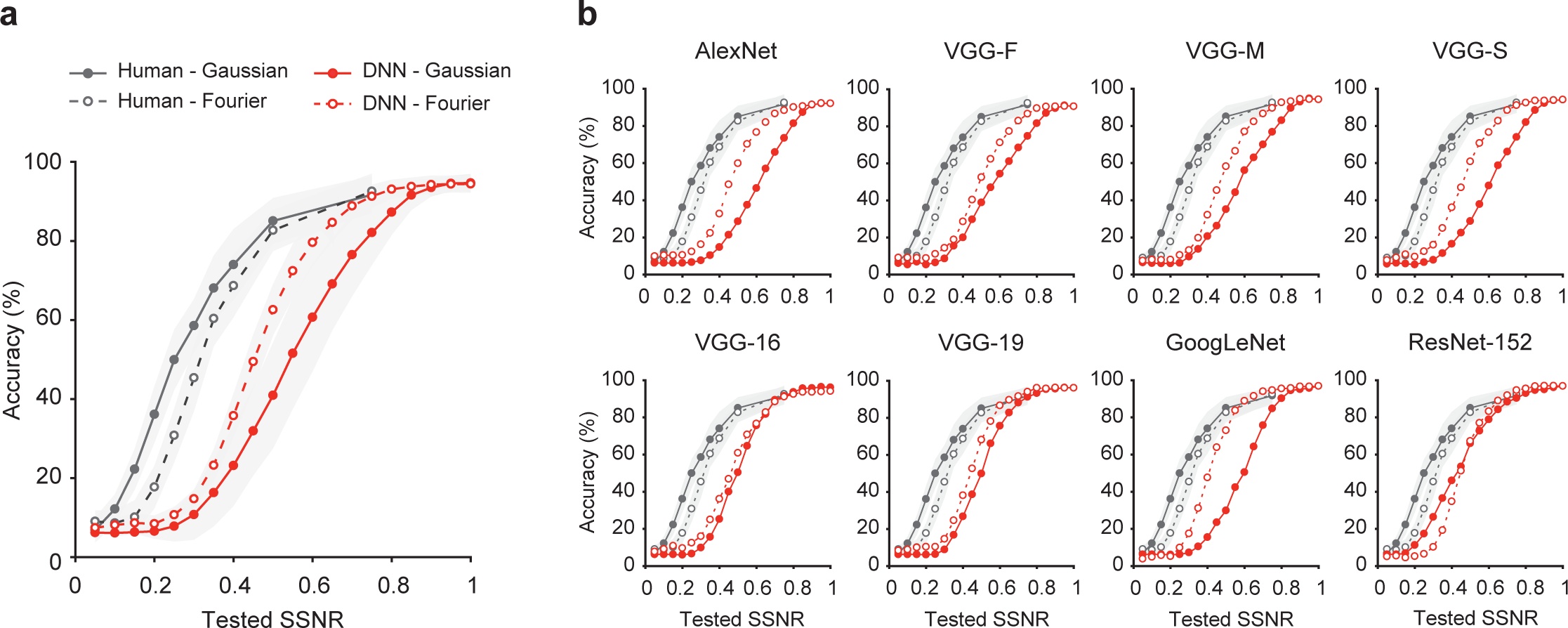
**a** Mean performance accuracy in a 16-alternative object classification task plotted as a function of SSNR level for human observers (black curves) and 8 standard pre-trained DNNs (red curves) with ± 1 standard deviation in performance indicated by the shaded area around each curve. Separate curves are plotted for pixelated Gaussian noise (solid lines with closed circles) and Fourier phase-scrambled noise (dashed lines with open circles). **b** Classification accuracy plotted as a function of SSNR level for individual pre-trained DNN models.

Of particular interest, the DNNs appeared to be impaired by noise in a manner that qualitatively differed from human performance. Spatially correlated noise proved more challenging to human observers, whereas the DNNs were more severely impaired by pixelated Gaussian noise (in 7 out of 8 cases). We fitted a logistic function to the performance accuracy data of each participant and each DNN to determine the threshold SSNR level at which performance reached 50% accuracy. This analysis confirmed that human observers exhibited much lower SSNR thresholds than DNNs, outperforming the DNNs by a highly significant margin at recognizing objects in pixelated noise (*t*(26) = 15.94, *p* < 10^-14^); they also outperformed DNNs at recognizing objects in Fourier phase-scrambled noise (*t*(26) = 12.29, *p* < 10^-11^). Moreover, humans showed significantly lower SSNR thresholds for objects in pixelated noise as compared to spatially correlated noise (0.255 vs. 0.315; *t*(19) = 13.41, *p* < 10^-10^), whereas DNNs showed higher SSNR thresholds for objects in pixelated noise as compared to spatially correlated noise (0.535 vs. 0.446; *t*(7) = 3.81, *p* = 0.0066).

The fact that spatially independent noise proved more disruptive for DNNs was unexpected, given that a simple spatial filtering mechanism, such as averaging over a local spatial window, should allow a recognition system to reduce the impact of spatially independent noise while preserving relevant information about the object. Instead, these DNNs are unable to effectively pool information over larger spatial regions in the presence of pixelated Gaussian noise.

We performed additional analyses to compare the patterns of errors made by DNNs and human observers, plotting confusion matrices for each of four SSNR levels (**Supplementary Figure 1**). Human performance remained quite robust even at SSNR levels as low as 0.2, as the majority of responses remained correct, falling along the main diagonal. Also, error responses were generally well distributed across the various categories, though there was some degree of clustering and greater confusability occurred among animate categories. In contrast, DNNs were severely impaired by pixelated noise when SSNR declined to 0.5 or lower, and showed a strong bias towards particular categories such as “hare”, “ cat” and “couch”. For objects in spatially correlated noise, the DNNs exhibited a preponderance of errors at SSNR levels of 0.3 and below, with bias towards “hare”, “owl” and “cat”.

### Development of a noise-training protocol to improve DNN robustness

We devised a noise-training protocol to determine whether it would be possible to improve the robustness of DNNs to noisy viewing conditions to better match human performance. For these computational investigations, we primarily worked with the VGG-19 network, as this pre-trained network performed quite favorably in comparison to much deeper networks (e.g., GoogLeNet, ResNet-152), and could be trained and evaluated in an efficient manner to evaluate a variety of manipulations.

First, we investigated the effect of training VGG-19 on images from the 16 object categories presented at a single SSNR level with either type of noise. After such training, the network was tested on a novel set of object images presented with the corresponding noise type across a full range of SSNR levels. We observed that training the DNN at a progressively lower SSNR level led to a consistent leftward shift of the recognition accuracy by SSNR curve (**Figure 3a**).

**Figure 3.**
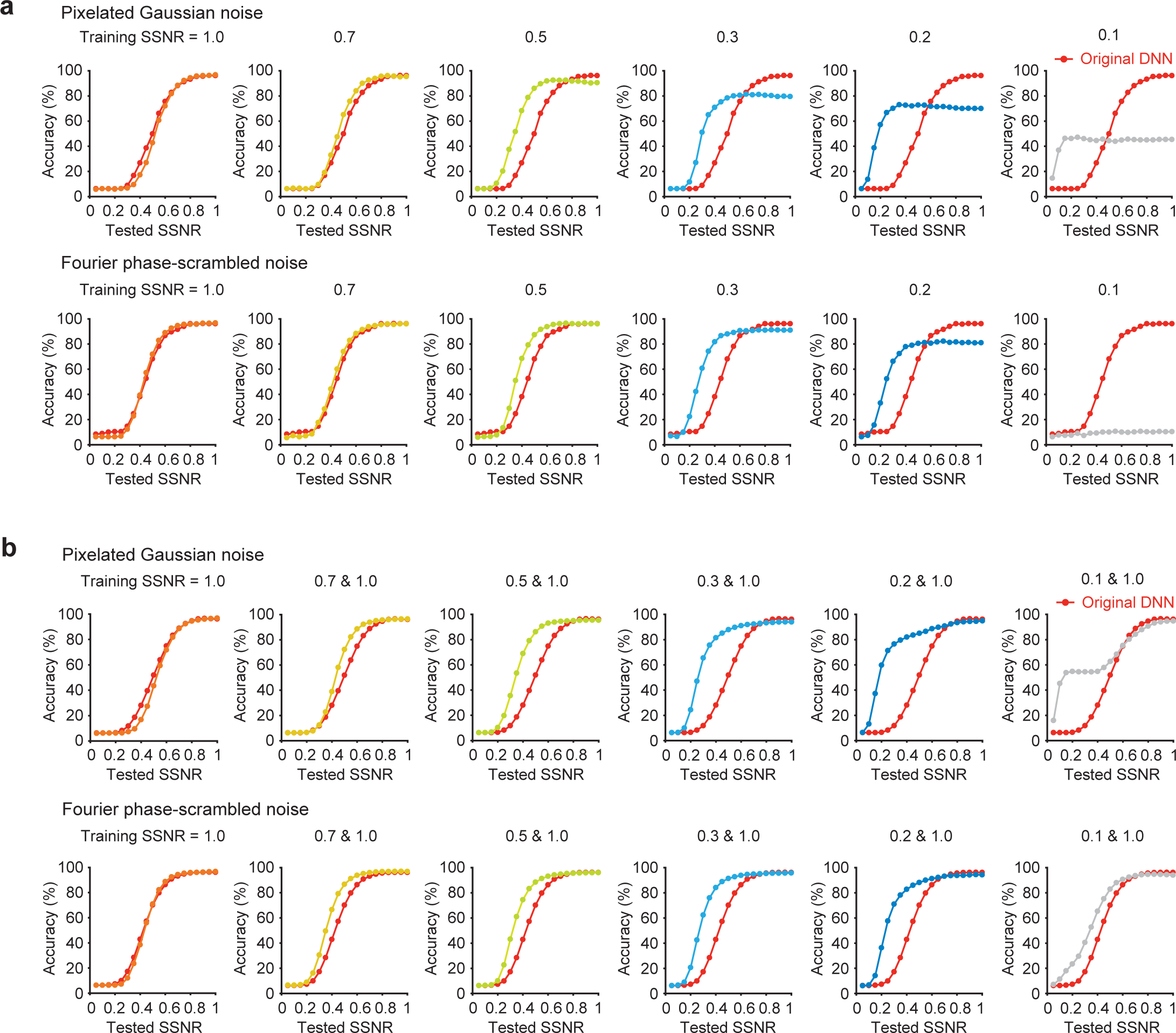
**a** Impact of training VGG-19 with object images presented at a single SSNR level (1.0, 0.7, 0.5, 0.3, 0.2, or 0.1) when evaluated with novel test images presented at multiple SSNR levels. Accuracy of pre-trained VGG-19 (red curve) serves as a reference in each plot. **b** Impact of training VGG-19 with a combination of noise-free images (SSNR 1.0) and noisy images at a specified SSNR level.

However, this improvement in performance for noisy images was accompanied by a loss of performance accuracy for noise-free images. The latter was evident from the prominent downward shift in the recognition accuracy by SSNR curve. Such loss of accuracy for noise-free images would be unacceptable for any practical applications of this noise-training procedure, and clearly deviated from human performance. Next, we investigated whether robust performance across a wide range of SSNR levels might be attained by providing intermixed training with both noise-free and noisy images. **Figure 3b** indicates that such combined training was highly successful, with the strongest improvement observed for noisy images presented at challenging SSNR level of 0.2. When the training SSNR was reduced to levels as low as 0.1, the task became too difficult and the learning process suffered.

Given the excellent performance of VGG-19 after training with images at 0.2 and 1.0 SSNR, we sought to compare noise-trained DNNs with human performance. **Figure 4a** shows that these noise-trained versions of VGG-19 performed far better than standard pre-trained DNNs at recognizing novel object images presented with the type of noise encountered during training. Moreover, noise training led to better performance for objects in pixelated Gaussian noise as compared to Fourier phase-scrambled noise, in a manner that better matched the qualitative performance of human observers. The noise-trained networks also seemed to outperform human observers on average.

**Figure 4.**
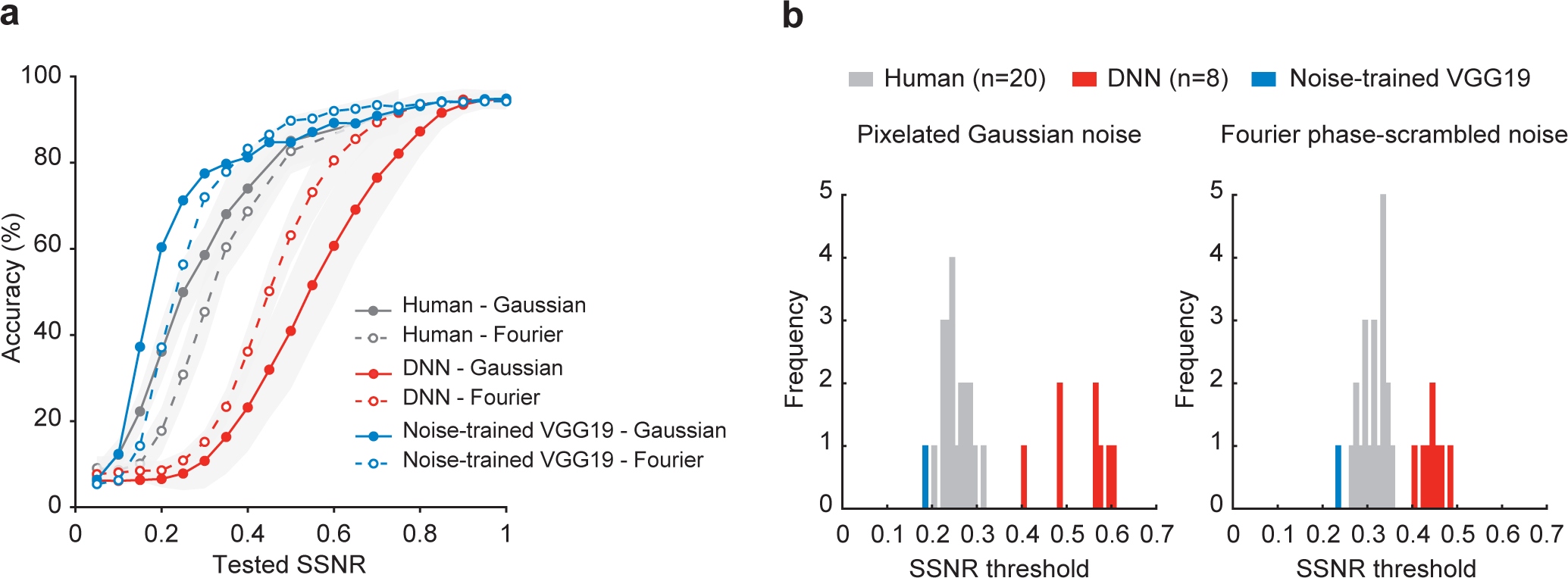
**a** Mean classification accuracy of noise-trained VGG-19 (blue), human observers (gray), and pre-trained DNNs (red) for objects in pixelated Gaussian noise (solid lines, closed circles) and Fourier phase-scrambled noise (dashed lines, open circles). **b** Frequency histograms comparing the SSNR thresholds of noise-trained VGG-19 (blue), individual human observers (gray), and 8 standard pre-trained DNNs (red).

To analyze these performance differences in detail, we fitted a logistic function to identify the SSNR thresholds of each DNN and human observer, separately for each noise condition. A histogram of SSNR thresholds revealed that both the Gaussian and Fourier noise-trained versions of VGG-19 outperformed all 20 human observers and all 8 original DNNs at recognizing objects in noise (**Figure 4b**). These results indicate that the noise-training protocol can greatly enhance the robustness of DNNs, such that they can match or surpass human performance when tasked to recognize objects in extreme levels of visual noise. These results are consistent with other recent reports [27]. However, such findings are insufficient to determine whether or not noise-trained DNNs have acquired visual representations that can account for the noise-robust nature of human vision.

### Image-level predictions of human behavioral performance

The ability to predict human recognition performance at the level of specific images constitutes one of the most stringent tests for evaluating DNN models; however, current DNN models have yet to adequately account for image-specific human performance [28]. Here, we devised a second behavioral experiment to evaluate whether noise-trained DNNs might be capable of predicting the noise threshold at which people can successfully recognize objects on an image-by-image basis.

Twenty observers were presented with each of 800 object images (50 per category), which slowly emerged from pixelated Gaussian noise. The SSNR level gradually increased from an initial value of 0 in small steps of 0.025 every 400ms, until the observer pressed a key to pause the dynamic display in order to make a categorization decision. A reward-based payment scheme provided greater reward for correct responses made at lower SSNR levels. After making a categorization response, participants used a mouse pointer to demarcate the portions of the image that they relied on for their recognition judgment. The resulting data allowed us to compare the similarity of humans and DNNs in their SSNR thresholds, as well as the portions of each image that were diagnostic for recognition judgments.

Mean performance accuracy was high (90.3%), and human SSNR thresholds for each image were calculated based on responses for correct trials only. Accordingly, SSNR thresholds were calculated for standard and noise-trained VGG-19 by requiring accuracy to reach 90%. Here, the standard DNN consisted of pre-trained VGG-19 that received an equal number of training examples from the 16 object categories using noise-free images only.

Although the standard DNN could predict human SSNR thresholds to some degree (see **Figure 5a**, slope = 0.32, *r* = 0.27, *t*(713) = 7.44, *p* < 10^-12^), the Gaussian noise-trained DNN provided a significantly better fit of human thresholds for individual images (slope = 0.67, *r* = 0.55, t(730) = 17.77, p < 10^-16^, comparison of noise-trained vs. standard DNN, *z* = 6.50, *p* < 10^-10^). These findings indicate that noise-trained DNNs provide a better model for predicting the critical noise level at which humans can recognize individual objects. That said, it should be noted that human-to-human similarity was greater still (mean *r* = 0.94, based on a split-half correlation analysis), indicating that further improvements can be made by future DNN models to account for human recognition performance.

**Figure 5.**
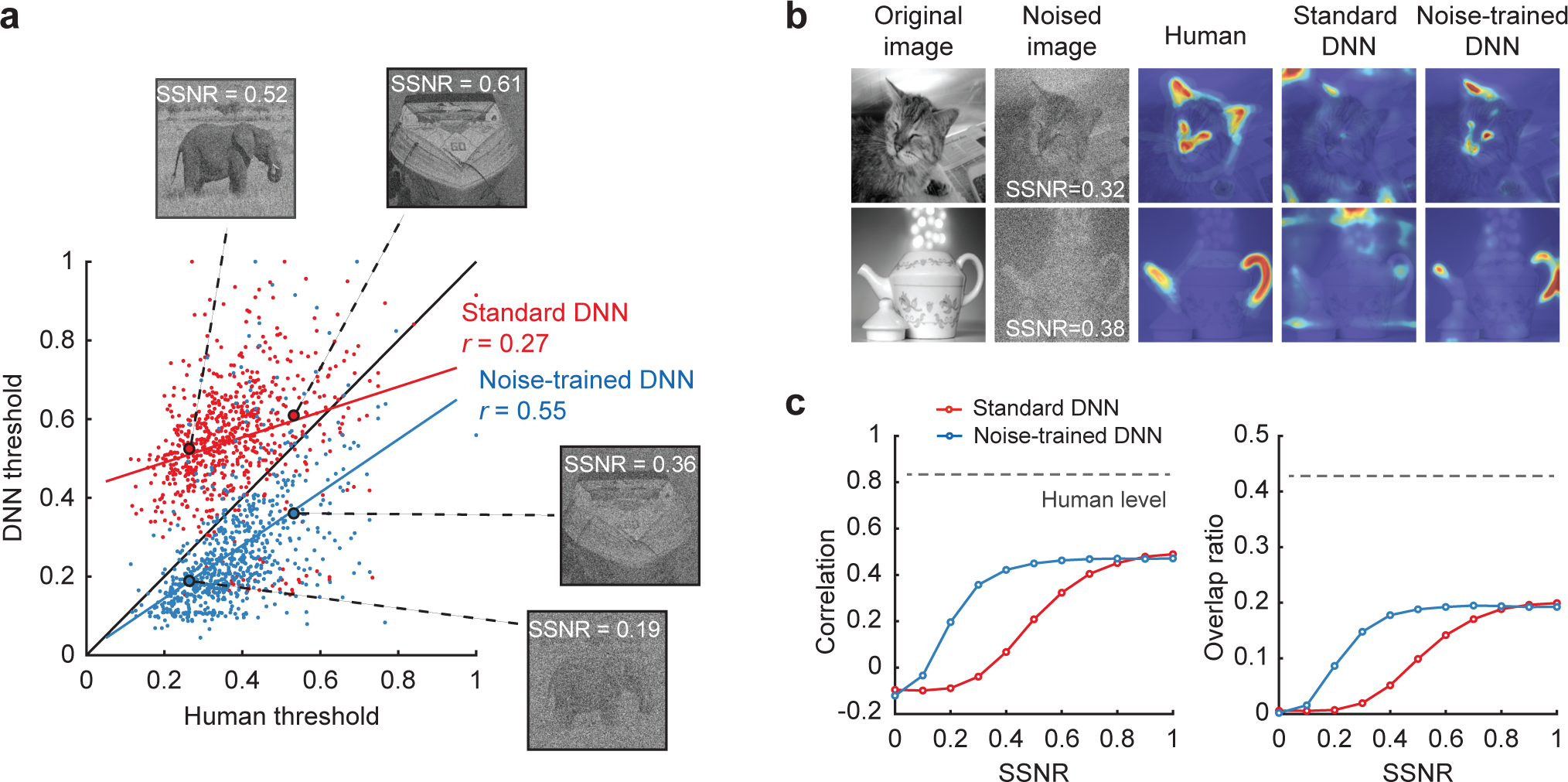
**a** Scatter plot comparing SSNR thresholds of human observers with the thresholds of standard VGG-19 (red) and noise-trained VGG-19 (blue). Each data point depicts SSNR thresholds for an individual object image. Examples of two object images, shown at the SSNR threshold obtained from standard or noise-trained networks. **b** Examples of diagnostic object features from human observers, standard VGG-19, and noise-trained VGG-19. The mean SSNR level at which human observers correctly recognized the objects is indicated. **c** Correlational similarity and overlap ratio of the spatial profile of diagnostic features reported by human observers and those measured in DNNs across a range of SSNR levels. Gray dashed lines indicate ceiling-level performance based on human-to-human correspondence.

To complement the diagnostic regions reported by human observers, we used layer-wise relevance propagation [40] to determine what portions of each image were important for the decisions of DNNs (see **Figure 5b**). We calculated the spatial correlation and amount of overlap between the diagnostic regions of humans and DNNs across a range of SSNR levels. Both standard and noise-trained DNNs performed quite well at predicting the diagnostic regions used by human observers at high SSNR levels of 0.8 or greater (**Figure 5c**). However, only the noise-trained DNN could reliably predict the diagnostic regions used by human observers in noisy viewing conditions. The above findings demonstrate that noise-trained DNNs can capture the fine-grained behavioral performance patterns of human observers when tasked to recognize objects in challenging noisy conditions.

### Characterizing network changes caused by noise training

Given that noise-trained DNNs provide an effective model for predicting human recognition of objects in noise, we sought to identify the stages of DNN processing that are most affected by training with a specific type of noise. We devised a layer-specific noise susceptibility analysis that required calculating the correlation strength between the layer-specific pattern of activity evoked by a noise-free image and the pattern of activity evoked by that same image when presented at varying SSNR levels (**Figure 6a**). Here, correlation strength should monotonically increase with increasing SSNR level (from an expected R value of 0 to 1.0), and the threshold SSNR level needed to reach a correlation value of 0.5 can then be identified. A lower threshold SSNR indicates greater robustness, whereas a higher threshold SSNR indicates greater noise susceptibility. We confirmed that the correlational similarity between layer-specific responses to a noise-free image and noisy image did indeed increase as a monotonic function of latter’s SSNR level (**Supplementary Figure 2**), and observed greater robustness in noise-trained than standard DNNs, especially in the higher layers.

**Figure 6.**
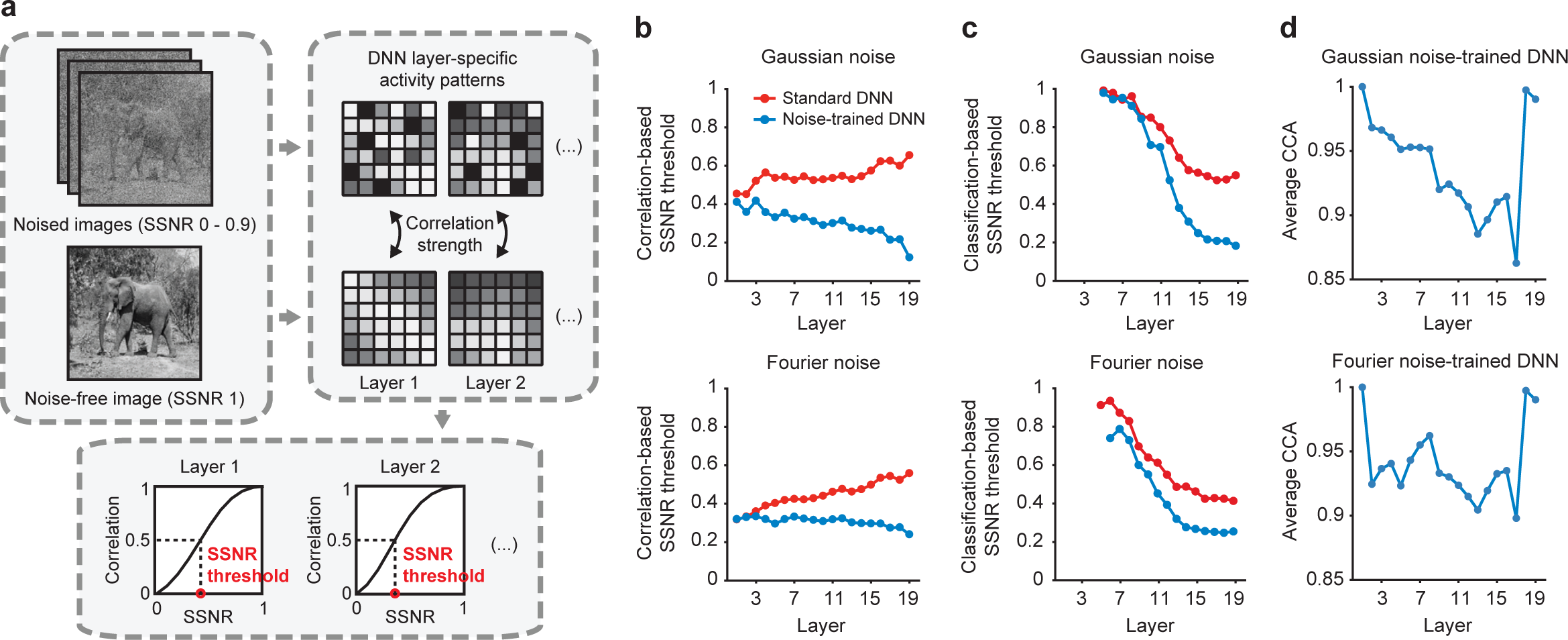
**a** Depiction of method used for layer-specific noise susceptibility analysis. **b** Correlation-based SSNR thresholds for pre-trained (red) and noise-trained (blue) versions of VGG-19 plotted by layer for objects shown in pixelated Gaussian noise or Fourier phase-scrambled noise. Higher SSNR thresholds indicate greater susceptibility to noise. **c** Classification-based SSNR thresholds plotted by layer for pre-trained and noise-trained networks. Multi-class support vector machines were used to predict object category from layer-specific activity patterns. **d** Similarity of feature representations for pre-trained and noise-trained versions of VGG-19, calculated using canonical correlation analysis (CCA).

As can be seen in **Figure 6b**, the standard and noise-trained DNNs exhibit quite similar SSNR thresholds in the first few layers but thereafter performance begins to diverge. For the standard DNNs, noise susceptibility gradually increases in progressively higher layers for both types of noise, implying that the contaminating effects of image noise tend to become amplified across successive stages of feedforward processing. After noise training, however, the network shows considerable improvement, especially in the middle and higher layers where the difference between standard and noise-trained networks most clearly diverges. For pixelated Gaussian noise, SSNR thresholds actually decrease across successive layers. In effect, the convolutional processing that occurs across successive stages of the noise-trained network leads to a type of de-noising process. This finding is consistent with the notion that the disruptive impact of spatially independent noise can be curtailed if signals over progressively larger spatial regions are pooled together in an appropriate manner to dampen the impact of random, spatially independent noise. This can be contrasted with the results for the DNN trained on objects in Fourier phase-scrambled noise. Here, the SSNR thresholds of the noise-trained network remain quite stable across successive layers, whereas the standard DNN becomes more susceptible to noise across successive layers.

As a complementary analysis, we measured classification-based SSNR thresholds by applying a multi-class support vector machine (SVM) classifier to the activity patterns of each layer of a given network. Each SVM was trained on activity patterns evoked by noise-free training images, and then tested on its ability to predict the object category of test stimuli presented at varying SSNR levels. The SSNR level at which classification accuracy reached 50% was identified as the classification-based SSNR threshold. For standard DNNs, we found that classification accuracy for noise-free test images gradually improved across successive layers of the network due to increased sensitivity to category information (**Supplementary Figure 3**), and this trend largely accounted the gradual improvement in classification-based SSNR threshold from one layer to the next (**Figure 6c**). Of greater interest, the divergence between standard and noise-trained networks became more pronounced in the middle and higher layers due to the benefits of noise training. These findings favor the notion that acquisition of noise robustness involves considerable modification of representations in the middle and higher layers of the noise-trained network. Consistent with this notion, a study of DNNs trained on auditory stimuli, specifically spoken words and music in the presence of real-world background noise, found that robustness to auditory noise was more pronounced in the higher layers of the DNN [41].

We further evaluated the extent to which the feature representations or weights in each layer changed as a consequence of noise training. This was done by performing canonical correlation between the weight matrices of the pre-trained DNN and the noise-trained DNN to assess their multivariate similarity. As can be seen in **Figure 6d**, training with Gaussian noise led to negligible change to the representations in layer 1, whereas progressively greater change was observed in the subsequent convolutional layers of the network. However, layers 18 and 19, which are fully connected and tend to represent more semantic rather than visual information, exhibited negligible change in their structured representations after noise training. For the DNN trained with Fourier phase-scrambled noise, we observed exhibited negligible changes in layer 1 and moderate changes in layers 2 through 17. Taken together, these analyses indicate that noise training leads to modifications to all convolutional layers of the DNN, with the exception of layer 1. Presumably, these changes account for the greater robustness to noise that was observed in our layer-wise measures of noise susceptibility (**Figure 6b,c**).

### Comparison of DNNs and human visual cortical responses to objects in noise

We conducted an fMRI study at 7 Tesla to measure human cortical responses to objects in noise and to assess their degree of correspondence with DNN object representations.

Observers were shown 16 object images (2 images from 8 selected categories) in each of 3 viewing conditions: without noise (SSNR 1.0), in pixelated Gaussian noise (SSNR 0.4) and in Fourier phase-scrambled noise (SSNR 0.4). During each image presentation, observers were instructed to perform an animate/inanimate discrimination task. Behavioral accuracy was high overall (97.1% for clean objects, 98.5% for Gaussian noise, 95.5% for Fourier phase-scrambled noise) and did not significantly differ between conditions (*F*(2, 21) = 1.27, *p* = .30).

First, we sought to determine whether the human visual cortex is more readily disrupted by spatially correlated noise than by spatially independent noise, as one might expect from our behavioral results from Experiment 1. We evaluated object discrimination performance of individual visual areas by training a multi-class SVM classifier on fMRI responses to clean object images and testing the classifier’s ability to predict the object category of both clean and noisy images using cross validation (see Methods). In early visual areas V1 through V4, object classification of cortical responses was most accurate for clean images, intermediate for objects in pixelated Gaussian noise, and poorest for objects in Fourier phase-scrambled noise (**Figure 7a**). Planned comparisons indicated that classification accuracy was significantly higher for clean objects than for objects in pixelated noise, and also higher for objects in pixelated noise as compared to Fourier phase-scrambled noise (*t*(7) > 4.7 in all cases, *p* < 0.0025). These fMRI results concur with the better behavioral performance that human observers exhibited for objects in pixelated noise as compared to Fourier phase-scrambled noise.

**Figure 7.**
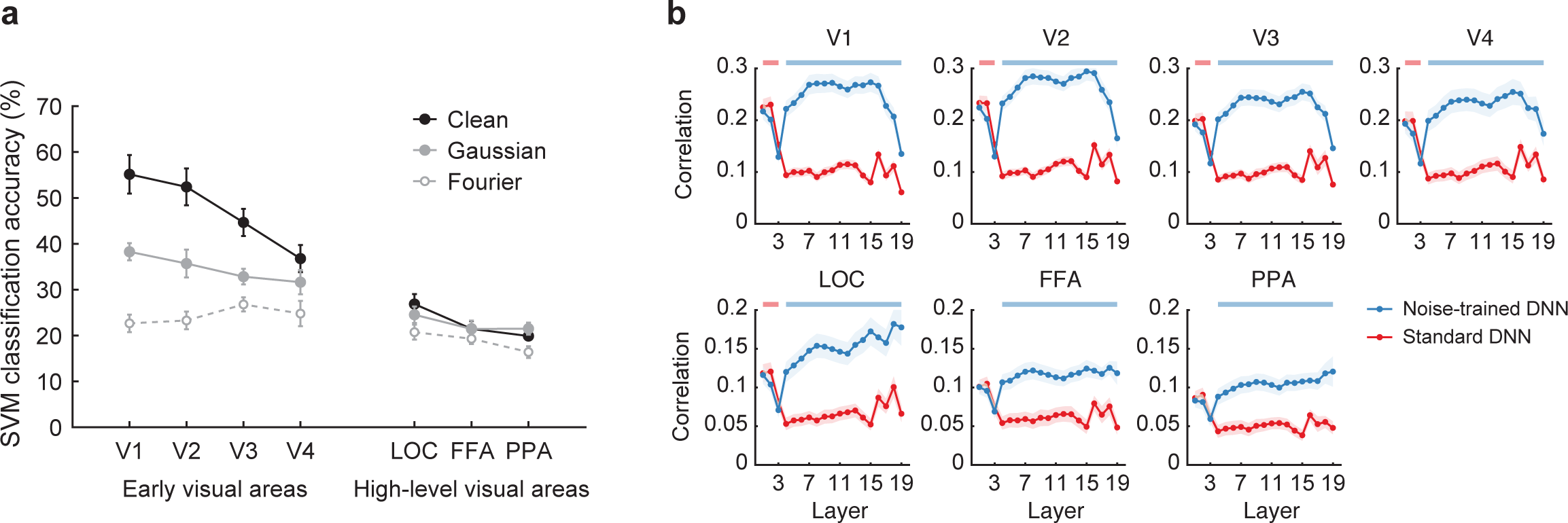
**a** Classification accuracy for fMRI responses in individual visual areas for clean objects (black filled circles), objects in pixelated Gaussian noise (gray filled circles) and Fourier phase-scrambled noise (gray open circles). Error bars indicate ±1 standard error of the mean (n = 8). Chance-level performance is 12.5%. **b** Correlational similarity of object representations obtained from human visual areas and individual layers of DNNs when comparing standard versus noise-trained networks (red vs. blue, respectively). Color-coded horizontal lines at the top of each plot indicate a statistically significant advantage (p < .01 uncorrected) for a given DNN at predicting human neural representations of the object images.

Classification accuracy for high-level object-sensitive areas was lower overall than was observed for early visual areas; this pattern of results is often found in studies of fMRI decoding. Interestingly, discrimination accuracy for objects in pixelated noise was not significantly different from that of clean objects in any of the high-level areas. These findings are consistent with the notion that greater spatial pooling of information by higher visual areas may attenuate the detrimental effects of spatially independent noise [32]. By contrast, classification accuracy was significantly better for objects in pixelated Gaussian noise as compared to Fourier phase-scrambled noise in the lateral occipital cortex (LOC, *t*(7) = 3.38, *p* < 0.025) and the parahippocampal place area (PPA, *t*(7) = 2.54, *p* < 0.05), though not in the fusiform face area (FFA, *t*(7) = 1.09, *p* = 0.31). Our findings indicate that object processing in both low- and high-level visual areas is more readily disrupted by spatially correlated noise than by spatially uncorrelated noise.

We evaluated the correspondence between human cortical responses and DNN representations by performing representational similarity analysis [13]. This involved calculating a correlation matrix of the responses to each of the 48 images (16 object images × 3 viewing conditions), separately for each visual area and for each layer of a DNN. After excluding the main diagonal, the resulting matrices reflected the similarity (or confusability) of responses to all possible pairs of object images. The similarity of the object representational spaces across humans and DNNs could then be determined by calculating the Pearson correlation between matrices obtained from human visual areas and DNNs. The noise-trained DNN consisted of VGG-19 trained on the 16 categories of objects presented with the both types of noise as well as noise-free images. The standard DNN consisted of pre-trained VGG-19 that received an equal number of training examples with noise-free images only.

**Figure 7b** shows the results of standard and noise-trained DNNs in terms of their layer-specific ability to predict the patterns of responses in human visual areas. In the lowest layers 1-3, standard DNNs exhibited a modest advantage over noise-trained DNNs. However, from convolutional layer 4 and above, noise-trained DNNs exhibited a clear advantage over standard DNNs at predicting the similarity structure of human cortical responses, while performing better overall. For early visual areas, the correspondence with noise-trained VGG-19 remained high throughout convolutional layers 4 through 16, and then exhibited a sharp decline in the fully connected layers 17-19. These fMRI results concur with the fact that the later fully connected layers of DNNs tend to represent more abstracted object information rather than visual information [14]. A different pattern of results was observed in high-level object-sensitive areas (LOC, FFA, PPA). Here, the correspondence with noise-trained DNNs remained high or tended to rise in the fully connected layers. Taken together, these results demonstrate that noise-trained DNNs provide an effective model to account for the pattern of visual cortical responses to objects in noise, whereas standard DNNs do not.

### Generalization of noise training to other DNNs and to other stimulus conditions

We conducted a series of studies with DNNs to assess whether noise training was effective at improving their performance across a wider range of conditions. Although the benefits of noise training were specific to the noise type encountered during training (**Supplementary Figure 4**), we found that it was possible to train a single DNN to acquire robustness to both pixelated Gaussian noise and Fourier phase-scrambled noise concurrently (**Supplementary Figure 5**). Likewise, we confirmed that other networks (e.g., ResNet-152) showed similar improvements in robustness after noise training with these 16 object categories (**Supplementary Figure 6**). We evaluated the impact of training VGG-19 on the full 1000-category image set from ImageNet with both types of noise, and found that the network was capable of recognizing objects in noise when discerning among a large number of categories (**Supplementary Figure 7**). These studies confirm that noise training is effective for much deeper networks and for large-scale image datasets. Further, they indicate that robustness to both spatially independent noise and spatially correlated noise can be learned concurrently by a single DNN.

We also investigated whether our noise-trained DNNs might show evidence of successful generalization to other types of image distortion, such as salt-and-pepper noise as well as low-pass and high-pass filtering. These analyses were motivated by a related study that reported extremely poor generalization whenever a DNN was trained on one type of image distortion and then tested with another [27]. In **Figure 8**, it can be seen that DNNs trained on pixelated Gaussian noise, or both Gaussian and Fourier phase-scrambled noise, can generalize well to object images corrupted by salt-and-pepper noise, outperforming standard pre-trained DNNs. By contrast, the DNN trained with Fourier phase-scrambled noise showed better performance at recognizing high-pass filtered images than the standard DNN, but with some cost in performance for low-pass filtered images. The DNN trained on both types of noise showed a similar improvement at recognizing high-pass filtered images, with no associated cost in recognizing low-pass filtered images. Thus, in contrast to earlier reports, we find that DNNs trained with noisy object images can generalize to other types of image distortions to some extent.

**Figure 8.**
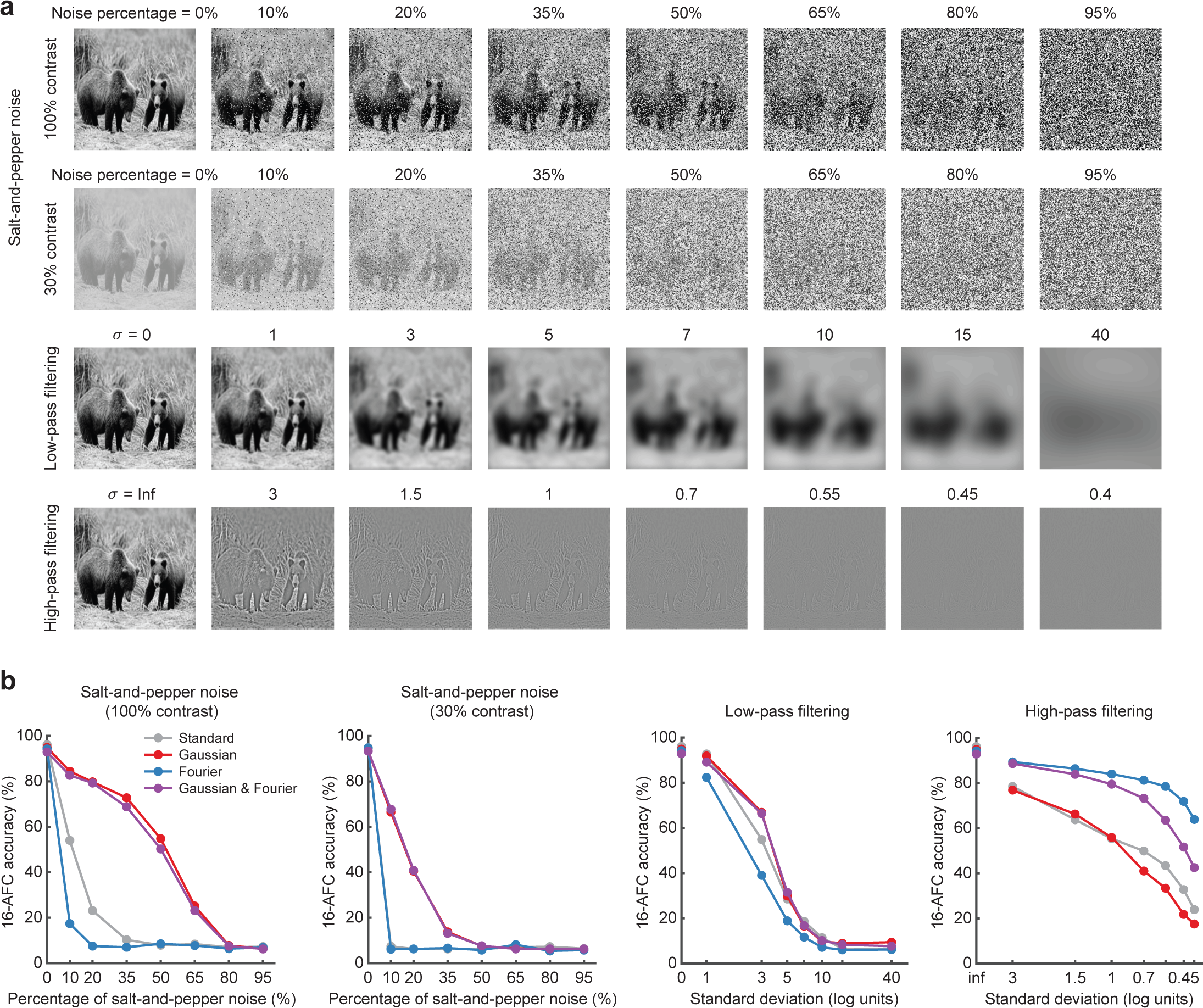
**a** Examples of images used to test the impact of salt-and-pepper noise, low-pass filtering and high-pass filtering on DNN performance. Image manipulations followed the methods described in [27]. **b** Performance accuracy of pre-trained and noise-trained versions of VGG-19 at recognizing images with different types and levels of image distortion.

Expanding on this question, we sought to ask whether DNNs, trained on objects in artificial noise, might show successful generalization to real world conditions of noise. Weather conditions such as rain, snow or fog are among the major causes of poor visibility for human observers. Using the 1000-category noise-trained VGG-19, we devised a test to determine whether noise training might have improved its ability to recognize novel real-world examples of objects in noisy weather. We focused on classification performance for 8 types of vehicles in the ImageNet data set. A web-based search protocol was used to gather candidate images of vehicles in noisy weather conditions, and test images were selected based on the ratings of three independent observers. The final test set consisted of 102 noisy vehicle images and 102 noise-free vehicle images (see **Supplementary Figure 8** for examples). **Figure 9a** shows that both standard and noise-trained versions of VGG-19 performed equally well at recognizing noise-free images of vehicles. By contrast, noise-trained VGG-19 outperformed the standard DNN at recognizing vehicles in noisy weather conditions. Performance was further analyzed according to the human-rated noise level of individual vehicle images. This analysis indicated that noise-trained VGG-19 performed significantly better with images rated as containing moderate or strong noise (**Figure 9b**). Thus, DNNs trained on images with artificially generated noise can successfully generalize, to some extent, to real-world examples of noisy viewing conditions, presumably due to shared statistical properties across real and artificial noise types.

**Figure 9.**
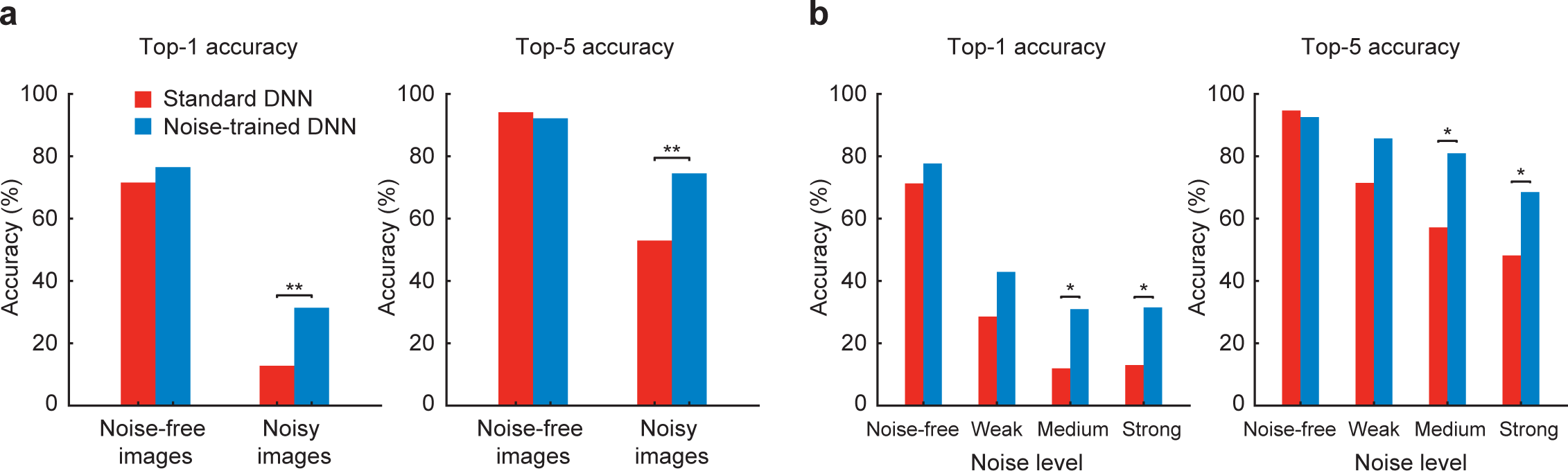
**a** Top1 and top5 accuracies of pre-trained VGG-19 (red) and noise-trained VGG-19 (blue) at classifying vehicles in noise-free or noisy weather conditions. Noise-trained VGG-19 outperformed pre-trained VGG-19 at recognizing noisy vehicle images (top1 accuracy, χ2 = 10.29, p = .0013; χ2 = 10.26, p = .0014). **b** Top1 and top5 accuracies sorted by noise-level rating. A statistical difference in performance was observed between models when the noise level was moderate or strong (χ2 > 4.5, p < .05 in all cases). Asterisks indicate * p < .05, ** p < .01.

## Discussion

Human vision is known to be robust across diverse contexts and viewing conditions. Here, we evaluated whether DNNs can provide a viable model of human visual processing when tested with challenge images consisting of objects embedded in randomly generated noise patterns. Our experiments revealed that well-known DNNs process visual information in a qualitatively different manner than human observers. They are disproportionately impaired by pixelated noise and are unable to spatially integrate relevant information in the presence of such noise. By contrast, noise-trained DNNs are more robust to spatially independent noise than to spatially correlated noise, consistent with our behavioral and fMRI results from human observers.

We sought to determine whether noise-trained DNNs can provide a suitable model to predict human behavioral and neural responses to individual object images. Our behavioral results revealed that noise-trained DNNs can reliably predict the threshold noise level at which individual images can be recognized by humans. Moreover, these noise-trained DNNs rely on similar diagnostic regions of images as human observers to make their decisions. fMRI data obtained from multiple levels of the human visual pathway revealed that the representational similarity structure of cortical responses to different object images bore a significant correspondence with the representational similarity of layer-specific responses in the noise-trained DNNs. Our findings indicate that noise-trained DNNs provide a suitable neural model for predicting both human behavior and neural responses to noisy, perceptually challenging images of objects.

One might ask what types of neural modifications are necessary to attain robustness to visual noise? A layer-specific analysis of the DNN models indicated that noise training led to widespread changes in the robustness of the network. The benefits of noise training tend to become magnified across successive stages of processing, presumably because the network learns to better leverage or readout relevant information from the lower layer while discounting irrelevant information associated with the noise sources. With respect to spatially independent noise, visual representations actually became more noise-robust across successive stages of processing, akin to a hierarchical denoising process, with the greatest benefit observed at higher levels of the network. These findings deviate from traditional notions of image processing, which typically rely on the modification of low-level visual filters to achieve noise filtering [42]. Here, we found negligible evidence of changes in noise robustness in the first few layers of the DNN. Instead, a divergence in performance between standard and noise-trained DNNs did not emerge until convolutional layer 4 (**Figure 6b**). Given that max-pooling occurs between convolutional layers 2 and 3, the visual representations in layer 4 would be more akin to complex cell than simple cell responses [6]. These findings imply that acquisition of noise robustness involves the modification of visual representations that occur after early stage filtering. In agreement with this DNN analysis, we found that the representational similarity of fMRI responses in the human visual cortex corresponded much better with noise-trained than standard DNNs, and that this advantage took place for noised-trained DNN representations in layer 4 and above.

Based on these findings, we speculate that robustness to visual noise is acquired, at least in part, through learning and experience. Consistent with this notion, research has shown that young children improve considerably in their ability to recognize objects in pixelated noise from ages 3 to 5, and that adults outperform children [43]. Studies of perceptual learning have further shown that adults can dramatically improve in their ability to discriminate faces in noise across training sessions, and that improvements in noise threshold are attributable to the learning of a more effective representation of the face stimuli [30]. Just as an experienced driver may feel more confident driving in rainy or snowy conditions, our experiences with suboptimal viewing conditions may have the unsought benefit of improving the robustness with which we see.

Previous studies of DNNs have shown that object classification can be disrupted by a small amount of non-random adversarial noise [44, 45]. More recent studies have shown that DNNs are also impaired by randomly generated pixelated noise, and can benefit from training with noisy object images [24, 25, 27]. For example, Geirhos et al. (2018) reported that DNNs can greatly improve after training with a particular type of image distortion or noise type, often exceeding human-level performance, albeit this study used the same object images for training and testing DNN performance. Despite these gains in performance, the researchers found that training with a particular type of noise, such as Gaussian noise, led to poor generalization to other types of noise, such as salt-and-pepper noise. By contrast, here we found that our Gaussian noise-trained network generalized quite well to salt-and-pepper noise. Also, training with Fourier phase-scrambled noise led to better performance for high-pass filtered images. Our findings are consistent with another recent report of improvements in generalization performance after training with noisy object images [46].

While these previous studies sought to measure the limitations of standard DNNs and the performance gains that can be achieved by training with noisy images, they did not directly address whether noise-trained DNNs might provide a suitable model of human behavioral performance, nor did they attempt to account for human neural data. By contrast, the goal of the present study was to determine whether DNNs can provide a suitable model to account for human behavioral and neural responses, especially when examined at a finer grain. We found that standard DNNs, commonly used in neuroscience research, process objects in noise in a qualitatively different manner than human observers. By contrast, noise-trained DNNs do indeed provide a viable model to account for the noise-robust nature of the human visual system.

It should be acknowledged that even though we could track the layer-wise changes in noise-trained DNNs and quantify their gains in noise robustness, a full understanding of how noise robustness is attained will require further research. Although DNNs are not black boxes and their computations are fully available to scrutiny and repeated interrogation, it remains a challenge for researchers to characterize how a series of non-linear computations performed by DNNs work to achieve a particular computational goal [23].

Although our study of DNNs was primarily focused on understanding biological vision, the noise training methods reported here may also be of relevance for applications in computer vision and artificial intelligence, including the development of autonomous vehicles, visually guided robots, and the analysis of real world images with adverse viewing conditions. For example, we found that DNNs can concurrently acquire robustness to both spatially independent noise and spatially correlated noise. Such noise-trained DNNs could perform better at recognizing objects in suboptimal viewing conditions, such as those caused by poor weather conditions, low light levels and/or sensor noise. Indeed, we found that noise-trained DNNs outperformed standard DNNs at recognizing vehicles in real-world noise conditions. Since it may be difficult to acquire large amounts of image data obtained under poor viewing conditions, the training of DNNs with high levels of artificial noise could prove useful as a method of data augmentation, to improve the robustness of DNN performance.

Finally, our study may contribute to future work by providing a paradigm for comparing human and DNN performance under stress-test visual conditions. Both human observers and DNNs perform at near-ceiling levels of accuracy with clean object images, limiting the ability to discern differences in performance. By contrast, objects in noise can be rendered progressively more challenging by reducing the SSNR level; this can allow one to obtain a sensitive measure of the critical SSNR threshold at which individual object images can be recognized. Given that we now know that standard DNNs are more impaired by spatially independent noise than correlated noise, unlike human observers, these measures can serve as a useful benchmark for evaluating whether future DNN models process visual information in a human-like manner. In particular, it will be of particular interest for future studies to investigate whether other types of DNN architectures, such as those that incorporate lateral or top-down connections [16, 47, 48], are capable of conferring greater robustness to DNNs, either with or without direct training on noisy images.

## Materials and Methods

### Participants

We recruited 23 participants in behavioral experiment 1 (18 females, 5 males), with 20 participants successfully completing both sessions of the study. A separate group of 23 participants were recruited in behavioral experiment 2 (14 females, 9 males), with 20 participants completing all 4 sessions of the study. Ages ranged from 19 to 33 years old.

An fMRI experiment was also carried out with a total of 11 participants (5 females), ages 21-49; data from 3 participants were excluded due to poor MR data quality. All participants reported having normal or corrected-to-normal visual acuity, and provided informed written consent using electronic consent forms (REDCap). The study was approved by the Institutional Review Board of Vanderbilt University (IRB #040945). Participants were compensated monetarily or through a combination of course credit and monetary payment.

### Visual stimuli

Object images were obtained from the ImageNet database [17], which is commonly used to train and test convolutional neural networks on object classification. We selected images from 16 categories for our experiments, which included a mixture of animate and inanimate object categories that would be recognizable to participants (**Figure 1b**). Both humans and DNNs were tested using images from the validation data set of ImageNet, with 50 images per category or 800 images in total. The test images were converted to grayscale to remove color cues that otherwise might boost the ability to recognize certain object categories in severe noise. DNNs were trained using images from the training set (1300 images per category), so the images used for testing were novel to both humans and DNNs.

In Experiment 1, objects were presented using two different types of visual noise: pixelated Gaussian noise and Fourier phase-scrambled noise (**Figure 1a**). To create each Gaussian noise image, the intensity of every pixel was randomly and independently drawn from a Gaussian distribution centered at 127.5, assuming that the range of possible pixel intensities (0 to 255) spanned ±3 standard deviations. For Fourier phase-scrambled noise, we calculated the average amplitude spectrum of the 800 images, generated a set of randomized phase values and performed the inverse Fourier transform to create each noise image. Such spatially correlated noise has some coherent structure that preserves the original power spectrum (close to a 1/F amplitude spectrum) but lacks strong co-aligned edges, due to the phase randomization, and can be described as having a cloud-like appearance. We avoided using the Fourier power spectrum of individual images to generate noise patterns, as residual category information could persist in this case, assuming that the categories differ to some extent in their overall power spectra.

To investigate the effect of noise on object visibility, we manipulated the proportion of object signal (*w*) contained in the object-plus-noise images. We describe the proportional weighting of this object information as the signal-to-signal-plus-noise ratio (SSNR), which has a lower bound of 0 when no object information is present (i.e., noise only) and an upper bound of 1 when the image consists of the source object only. SSNR differs from the more conventional measure of signal-to-noise ratio (SNR), which has no upper bound. Given a source object image defined by matrix ***S*** and a noise image ***N***, we can create a target image ***T*** with SSNR level of *w* as follows:

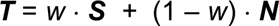

After the contrast-adjusted original image and the noise pattern were summed, any intensity values that fell beyond the 0-255 range were clipped. Clipping was modest as the standard deviation of the Gaussian noise distribution was 255/6.

### Behavioral experiment 1

Participants were tested with either pixelated Gaussian noise or Fourier phase-scrambled noise, in two separate behavioral sessions. To control for order effects, half of the participants were presented with pixelated Gaussian noise in the first session and while the other half were first presented with Fourier phase-scrambled noise.

In each session, participants were briefly presented with each of 800 object images for 200ms at a specified SSNR level, and had to make a 16-alternative categorization response thereafter using a keyboard. Noisy object images were presented at 10 possible SSNR levels (0.05, 0.1, 0.15, 0.2, 0.25, 0.3, 0.35, 0.4, 0.5, and 0.75). The highest SSNR level was informed by a pilot study that indicated that human accuracy reached ceiling levels of performance by an SSNR level of 0.75. Five images per category were assigned to each SSNR level, and image assignment across SSNR levels was counterbalanced across participants. The order of image presentation was randomized. The experiment was implemented using MATLAB and the Psychophysics Toolbox (http://psychtoolbox.org/).

### Behavioral experiment 2

This study measured participants’ SSNR thresholds for each of 800 object images over a series of 4 behavioral sessions. For this experiment, only pixelated Gaussian noise was evaluated. On each trial, a single noise image was generated and combined with a source object image, and the target image gradually increased in SSNR level by 0.025 every 400ms, until the participant felt confident enough to press a key on a number pad to halt the image sequence and then make a 16-alternative categorization response. Next, participants used a mouse pointer to “paint” the portions of the image that they found to be most informative for their recognition response.

After each trial, participants received visual feedback, based on a point scheme designed to encourage both fast and accurate responses. For correct responses, up to 200 points could be earned at the beginning of the image sequence (SSNR = 0), and this amount decreased with increasing SSNR, dropping to just 6 points at an SSNR level of 1. Incorrect responses were assigned 0 points. The participants received monetary payment scaled according to the total number of points earned across the 4 sessions.

### MRI scanning parameters

MRI data were collected using a 7-Tesla Philips Achieva scanner with a 32-channel head coil at the Vanderbilt University Institute for Imaging Science. We collected fMRI data using single-shot T2*-weighted gradient echo echo-planar imaging at a 2mm isotropic voxel resolution (TR 2s; TE 25ms; flip angle 63°; SENSE acceleration factor 2.9, FOV 224×224 mm; 46 slices with no gap; phase-encoding in AP direction). To mitigate image distortions caused by inhomogeneity, an image-based shimming technique was used. A T1-weighted 3D-MPRAGE anatomical scan was collected in the same session at 1mm isotropic resolution. Separately, retinotopic data were acquired using a 3-Telsa Philips Intera Achieva MRI scanner equipped with a 32-channel head coil, with fMRI data acquired at 3mm isotropic resolution (TR 2s; TE 35ms; flip angle 80°; FOV 240×240 mm; 36 slices).

### fMRI experiment

For the fMRI experiment, we selected 16 object images to characterize neural response patterns to objects in visual noise. The images included 2 examples drawn from 8 categories (bear, bison, elephant, hare, jeep, sports car, table lamp, teapot) whose difficulty levels were closely matched based on the reported SSNR levels from Experiment 2. Each object image was presented noise-free, embedded in pixelated Gaussian noise, and embedded in Fourier phase-scrambled noise. For the noise conditions, we chose an SSNR level of 0.4 as human performance dropped significantly by this noise level but was still accurate enough to be expected to lead to reliable neural responses. To control for potential order effects, the images were divided into two sets. In the first half of the experiment, one set was presented noise-free while the other set of object images appeared in each of the two types of noise. In the second half of the experiment, the assignment to noisy and noise-free conditions was reversed. Across participants, we counterbalanced how the objects were assigned to noisy and noise-free conditions across the two halves of the experiment. Each fMRI run consisted of 8 clean images, 8 Gaussian noise images, and 8 images of objects in Fourier phase-scrambled noise, presented in randomized order. On average, participants performed a total of 10 experimental runs with each image shown 5 times for a given condition.

Participants were instructed to maintain fixation on a central fixation point throughout each experimental run and to report whether each presented image was animate or inanimate using an MRI-compatible button box in the scanner. Each image from a stimulus set was centrally presented in a 9 × 9° window for 4 seconds, flashing on and off every 250ms, and followed by a 6-second fixation rest period. The order of the 24 images was randomized every run, and each run lasted approximately 4.4 minutes. We additionally ran 2 runs of a functional localizer, in which participants viewed blocked presentations of grayscale images of faces, objects, houses, and scrambled objects. A subset of the participants were scanned on a separate day for retinotopic mapping which used a standard phase-encoded measurement with rotating wedges and expanding rings [49].

### fMRI data preprocessing and analysis

Data were preprocessed and analyzed using FSL, Freesurfer, and custom MATLAB scripts. The following standard preprocessing was applied: motion correction using MCFLIRT [50], slice-time correction, and high-pass temporal filtering with a cutoff frequency of 0.01 Hz. No spatial smoothing was applied. Functional images were then registered to each participant’s 3D- MPRAGE anatomical scan using Freesurfer’s bbregister [51].

Boundaries between early visual areas V1-V4 were manually delineated from a separate retinotopic mapping session, using FSL and Freesurfer software. For those who did not perform retinotopic scanning, areas V1-V4 were predicted from the anatomically defined retinotopy template [52]. A general linear model analysis was used to identify visually responsive voxels corresponding to the stimulus location, as well as category-selective voxels.

In conjunction with the retinotopic maps, a statistical map of the stimulus versus rest contrast of our functional localizer was used to define functionally active voxels in V1-V4 using a threshold of t > 7 uncorrected. The fusiform face area (FFA) was identified by contrasting faces versus all other stimulus conditions (objects, houses, scrambled stimuli) and identifying voxels in the fusiform gyrus that exceeded a threshold of t > 3 uncorrected. Similarly, the parahippocampal place area (PPA) consisted of voxels in the parahippocampal gyrus that responded more strongly to houses than to all other stimulus conditions (t > 3 uncorrected). Finally, the lateral occipital cortex (LOC) was defined by contrasting objects versus scrambled objects (t > 3 uncorrected).

Each voxel’s time series was first converted to percent signal change, relative to the mean intensity across the run, and the averaged response of TRs 3 to 5 post-stimulus onset was used to estimate stimulus responses. Response amplitudes to each stimulus were then normalized by run to obtain an overall mean of 0 and standard deviation of 1. For each visual area, the multivariate response pattern to a given stimulus was converted into a data vector with associated category label, to be used for training or testing a classifier. We trained a multi-class linear SVM classifier to predict the object category of each stimulus, separately for each region of interest and viewing condition, using the LIBSVM MATLAB toolbox with the default parameter settings [53]. The trained SVM was then tested on independent test runs, using a leave-one-run-out cross-validation procedure [7]. (Note that the object category decoding analysis is sensitive to consistency of fMRI responses at both the image level and category level, and we confirmed that essentially the same pattern of classification results was found in early visual areas when decoding was performed to predict the specific image.) We required that classification accuracy for V1, the most reliable visual area for decoding, exceed a minimum of 20% (chance level 12.5%) when averaged across all 3 viewing conditions; otherwise, the data from that participant were excluded due to poor reliability. Data from three participants were excluded based on these criteria, and reported results are based on the data of 8 participants.

To compare the representations of DNNs to those in the human visual cortex, we analyzed the responses of all units within each layer of the DNN to each object image. The responses of a given unit to the set of object images were normalized and converted to z-scores. Next, we calculated the correlational similarity of the responses to all possible pairs of images by computing a 48 × 48 correlation matrix. After setting the main diagonal values to 0, the remaining values solely reflected the correlational similarity of responses to different object images for that layer. The representational structure of these object responses of the DNN could then be compared to the representational structure of object responses obtained from human visual areas by calculating the Pearson correlation coefficient between the correlation matrices. For statistical testing, the Fisher z-transform was applied to these correlation values obtained from each participant when comparing a visual area to a specific layer of a DNN, and t-tests were used to test for significant differences between Pearson correlation values.

### Deep neural networks

We evaluated the performance of 8 pre-trained convolutional neural networks (CNNs) using the MatConvNet toolbox [54]: AlexNet, VGG-F, VGG-M, VGG-S, VGG-16, VGG-19, GoogLeNet, and ResNet-152 [36–39]. All networks were pre-trained on the ImageNet 1000-category classification task. Performance on the 16-category classification task was evaluated by determining which of the 16 categories had the highest softmax response to a given image.

The training of CNNs with noisy object images was primarily performed using MatConvNet (version 1.0-beta25), with ancillary analyses performed using PyTorch (version 1.6.0). The majority of noise training experiments were performed using VGG-19, although we also confirmed that similar benefits of noise training were observed for AlexNet and ResNet-152.

For 16-category training, all DNNs were trained using stochastic gradient descent over a period of 20 epochs with a fixed learning rate of 0.001, batch size of 24, weight decay of 0.0005, and momentum of 0.9. All weights in all layers of the network were initialized from pre-trained models and were allowed to vary during the training, using backpropagation of the multinomial logistic loss across all 1000 classes. For our first set of analyses, pre-trained VGG-19 was trained with noisy object images presented at a single SSNR level (**Figure 3a**), using images from the 16 categories in the ImageNet training set (20,800 images in total). Separate networks were trained with either pixelated Gaussian noise or Fourier phase-scrambled noise. Training at a single SSNR level led to better performance for noisy object images but poorer performance for noise-free objects. Subsequently, we trained VGG-19 using a combination of noise-free and noisy images, typically using an SSNR level of 0.2 for most experiments. The VGG-19 model used to approximate human SSNR thresholds in Experiment 2 was trained with objects in pixelated Gaussian noise across a full range of SSNR levels from 0.2 to 1. The standard DNN used to fit human SSNR thresholds consisted of pre-trained VGG-19 that received the same number of training examples from the 16 categories using noise-free images only.

For training examples, we used the standard data augmentation pipeline provided by MatConvNet. Training images were derived from the original images by randomly cropping a rectangular region (with width-to-height aspect ratios that randomly varied from 66.67% to 150%) that subtended 87.5% of the length of the original image. The cropped image was resized to 224 × 224 pixels to fit most of the DNN models (except for AlexNet, which used 227 × 227 pixels). Additionally, the intensity of each of the RGB channels was shifted by a small offset, randomly sampled from a Gaussian distribution with a standard deviation of about 3. The images were then converted to grayscale. Finally, after the SSNR manipulation was applied (as described in ***Visual stimuli***), the average pixel intensity across training samples was calculated and subtracted from each training image.

We trained a 1000-category version of VGG-19 with the full set of training images from ImageNet; these were presented either noise-free, with pixelated Gaussian noise (SSNR 0.2) or with Fourier phase-scrambled noise (SSNR 0.2). Color information from these images was preserved but the same achromatic noise pattern was added to all 3 RGB channels for noise training. The network was trained over 10 epochs using a batch size of 64. All other training parameters were the same as those used in training the 16-category-trained VGG-19.

We quantified the accuracy of standard and noise-trained DNNs at each of 20 SSNR levels (0.05, 0.1, 0.15, … 1). Unlike the human behavioral experiments, DNN performance could be repeatedly evaluated tested without concerns about potential effects of learning, as network weights were frozen during the test phase. The DNN was presented with all 800 object test images at every SSNR level to calculate the accuracy by SSNR performance curve. A 4- parameter logistic function was fitted to the accuracy by SSNR curve and the SSNR level at which accuracy reached 50% was identified as the SSNR threshold for Experiment 1.

For the layer-specific noise susceptibility analysis, we evaluated the stability of the activity patterns evoked by objects presented in progressively greater levels of noise, by calculating the Pearson correlation coefficient between responses to each noise-free test image and to that same image presented at varying SSNR levels. Analyses were performed on each convolutional layer after rectification, the fully connected layers and the softmax layer of VGG-19. A logistic function was fitted to the correlation by SSNR data for each layer, and the SSNR level at which the correlation strength reached 0.5 was identified as the SSNR threshold. If some positive correlation was still observed when SSNR level was 0, then the range of correlation values were linearly rescaled to span a range of 0 to 1, prior to calculating the SSNR threshold.

For the layer-specific classification analysis, multi-class support vector machines (SVM) were trained on the activity patterns evoked by noise-free objects from each of the 16 categories, using data obtained from individual layers of the DNN. After training, the SVMs were tested using the 800 novel test images presented at varying SSNR levels. The SSNR level at which classification accuracy reached 50% (chance level performance, 1/16 or 6.25%) was identified by fitting a logistic function, and served as the classification-based SSNR threshold.

### Layer-wise relevance propagation

Layer-wise relevance propagation is a method that identifies diagnostic features that contribute to the prediction of a network [40]. To do so, the method decomposes the network’s output with respect to contributions of individual units, termed relevance scores *R* as defined below, and back-propagated the scores to the input layer:

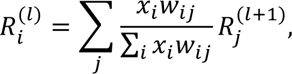

where *R*_*i*_^(*l*)^ is the relevance score of the unit *i* at layer *l*, *x*_*i*_ is the response of the unit *i* at layer *l*, and *w*_*i,j*_ is the weight connecting the unit *i* at layer *l* to unit 0 at layer *j*+1. Layer-wise relevance propagation differs from other gradient-based methods in that it takes into account both gradients and unit activations, and may thereby better capture the set of features that are responsible for the network’s classification response. In addition to the original implementation (i.e., LRP-0), several variants have been suggested including LRP-ε, LRP-γ, and LRP-zβ [55]. Following the guidance of Montavon et al. (2019), we implemented a VGG19-based custom PyTorch script as follows: LRP-0 from the 15^th^ to 19^th^ layers, LRP-ε (ε = 0.25) from the 9^th^ to 14^th^ layers, LRP-γ (γ = 0.05) from the 2^nd^ and 8^th^ layers, and LRP-zβ (lower bound = -1.99 and upper bound = 2.44) for the 1^st^ layer. To create pixel-wise heatmaps, the relevance scores in the pixel space were summed over the rgb channels. Only positive values were taken into account in order to focus on the category-relevant features of a selected object.

### Data availability

The experimental code, code for noise training of DNNs, as well as the human behavioral and neural data will be made available on open science framework upon publication of this work.

## Acknowledgements

The authors would like to thank Malerie McDowell, Echo Sun, Feyisayo Adegboye and Haley Frey for technical assistance.

## Author Contributions

FT devised the study, experimental design, protocol for noise training of DNNs, layer-specific noise susceptibility analysis, and provided guidance on data analysis. DM coded the first behavioral experiment; HJ coded the second behavioral experiment and the fMRI experiment. HJ coded the DNN networks, conducted all of the data analyses, and devised the layer-specific classification analysis. FT and HJ wrote the paper.

