## Supplementary Figures for "Noise-robust recognition of objects by humans and deep neural networks"

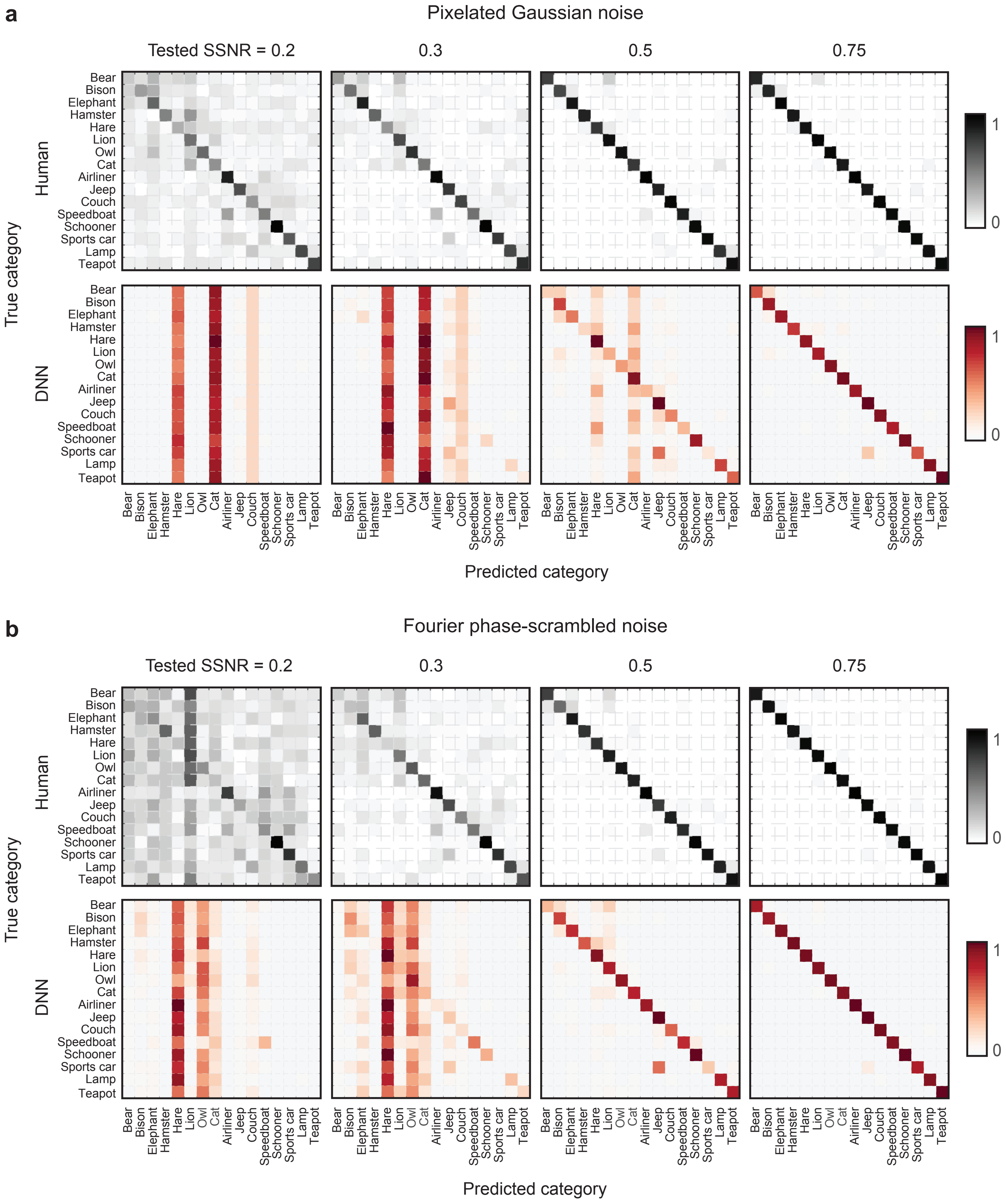


**Supplementary Figure 1.** Confusion matrices of human observers and 8 standard pre-trained DNNs. Plots show the relative frequency of predicted category responses (columns) with true categories organized by rows. Confusion matrices are provided for 4 SSNR levels, separately for objects in pixelated Gaussian noise **(a)** and Fourier phase-scrambled noise **(b)**.

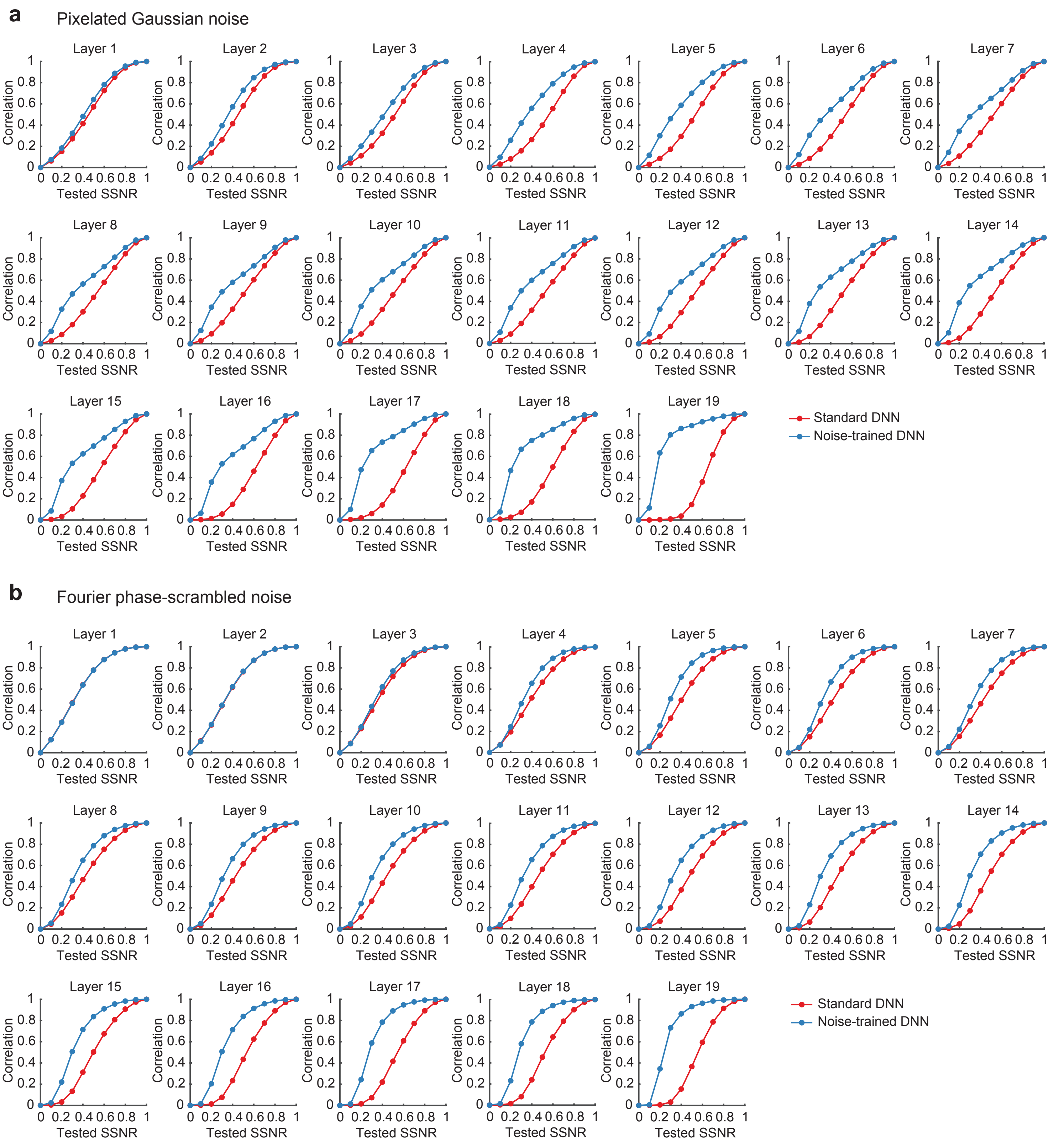


**Supplementary Figure 2.** Correlational similarity of layer-specific response patterns to noise-free objects and the same objects presented at varying SSNR levels. Results are plotted by layer for pre-trained VGG-19 (red) and noise-trained VGG-19 after training and testing with either pixelated Gaussian noise **(a)** or Fourier phase-scrambled noise **(b)**.

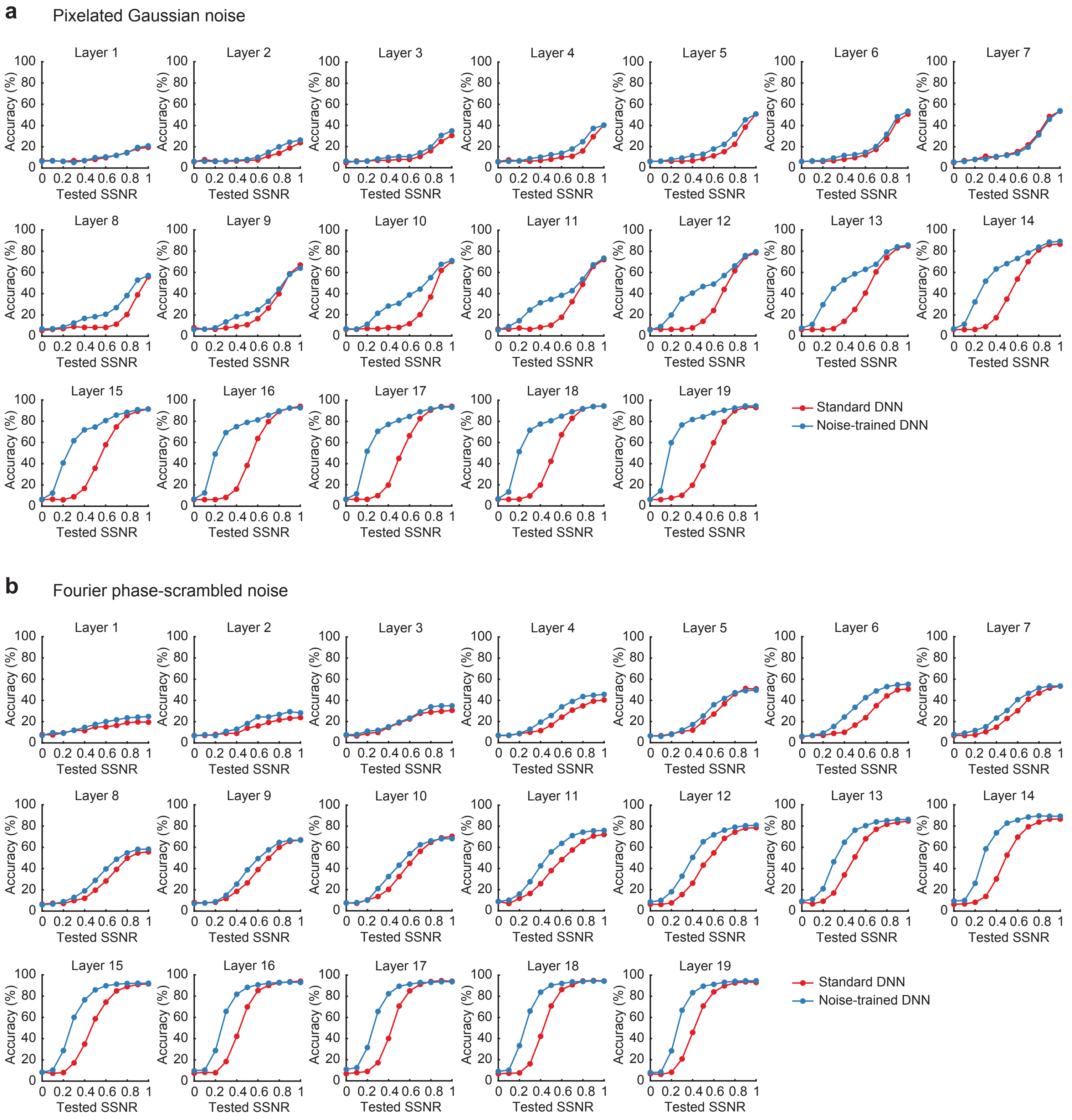


**Supplementary Figure 3.** Classification accuracy for SVMs tested on activity patterns obtained from specific layers of pre-trained or noise-trained VGG-19. SVMs were initially trained on activity patterns evoked by noise-free images. Subsequently, they were tested with novel test images in pixelated Gaussian noise **(a)** or Fourier phase-scrambled noise **(b)** at varying SSNR levels.

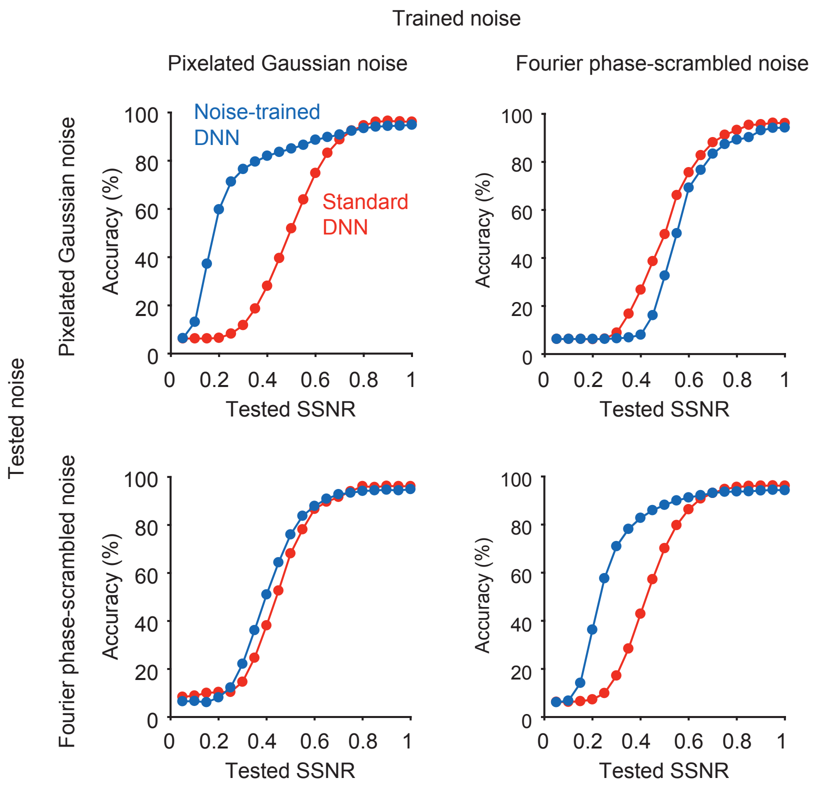


**Supplementary Figure 4.** Mean classification accuracy of noise-trained VGG-19 (blue) when trained with objects in either pixelated or Fourier phase-scrambled noise, and subsequently tested on either type of noise. Performance of pre-trained VGG-19 (red), which lacked noisy image training, is provided for comparison.

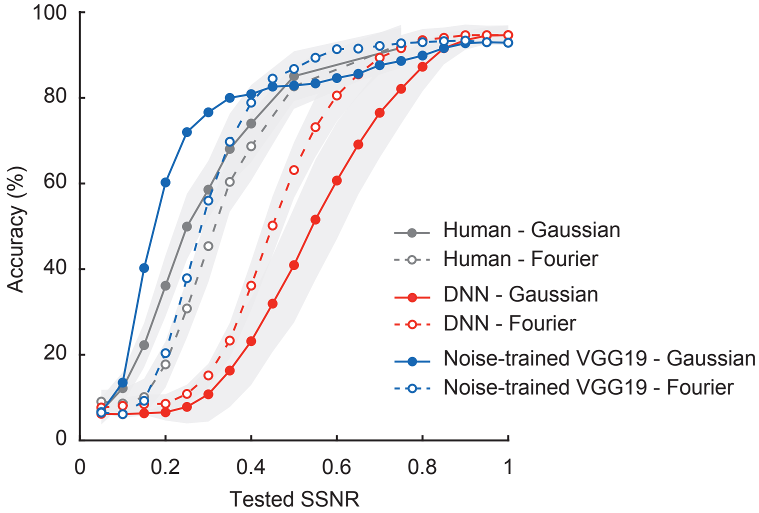


**Supplementary Figure 5.** Mean classification accuracy of noise-trained VGG-19 (blue) after training on a combination of noise-free images from the 16 categories, objects in pixelated noise (SSNR 0.2), and objects in Fourier phase-scrambled noise (SSNR 0.2). Performance of pre-trained DNNs and human observers is included for comparison, as in Figure 4.

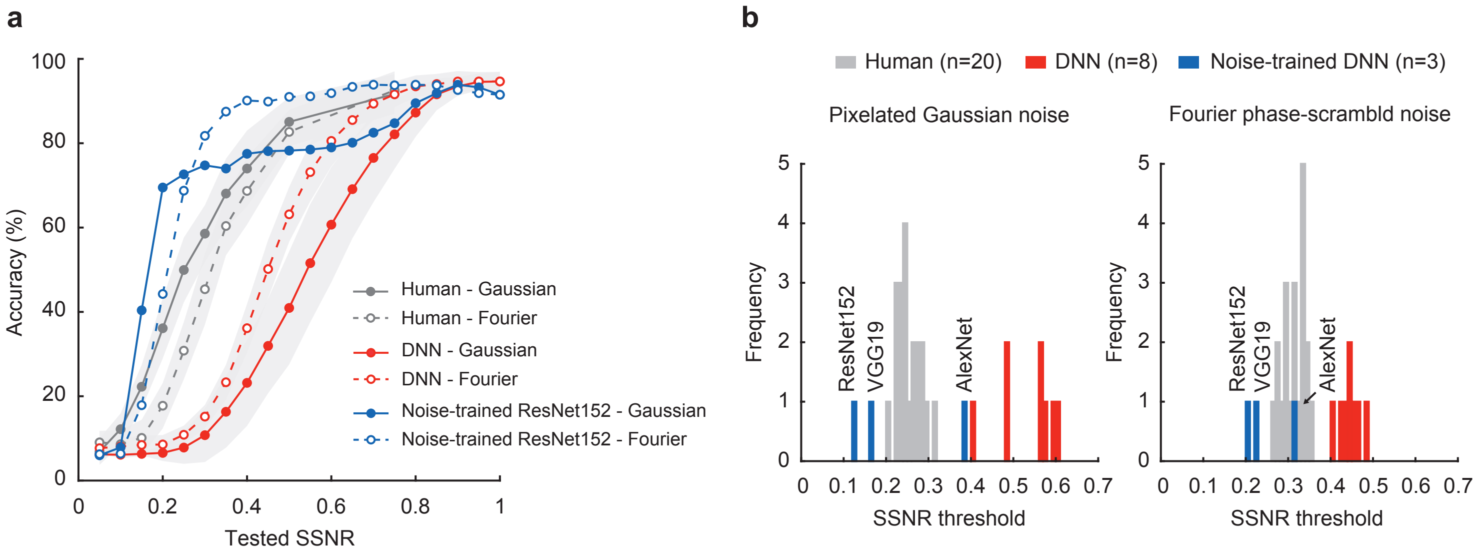


**Supplementary Figure 6.** a Mean classification accuracy of noise-trained ResNet-152 (blue), human observers (gray), and pre-trained DNNs (red). Here, noise-trained ResNet-152 was trained on noise-free images and objects in both types of noise (SSNR 0.2). **b** Frequency histograms of SSNR thresholds to attain 50% recognition accuracy for human observers, pre-trained DNNs, and DNNs trained with both types of noise plus noise-free images (i.e., VGG-19, ResNet-152 and AlexNet).

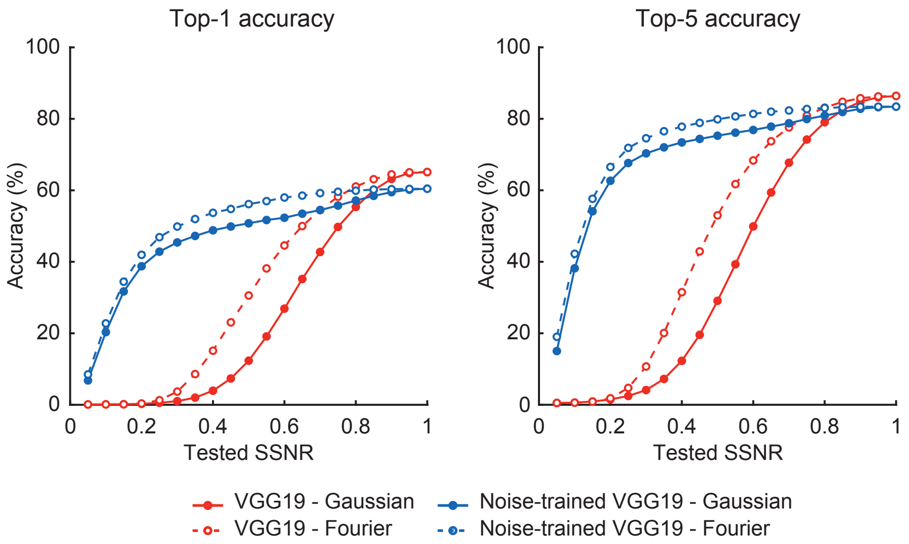


**Supplementary Figure 7.** Top1 and top5 accuracies of pre-trained VGG-19 (red) and noise-trained VGG-19 (blue) on the 1000-category ImageNet classification task. Here, the noise-trained VGG-19 was trained on ImageNet 1,000 categories with noise-free images and noisy images presented at 0.2 SSNR from both types of noise.

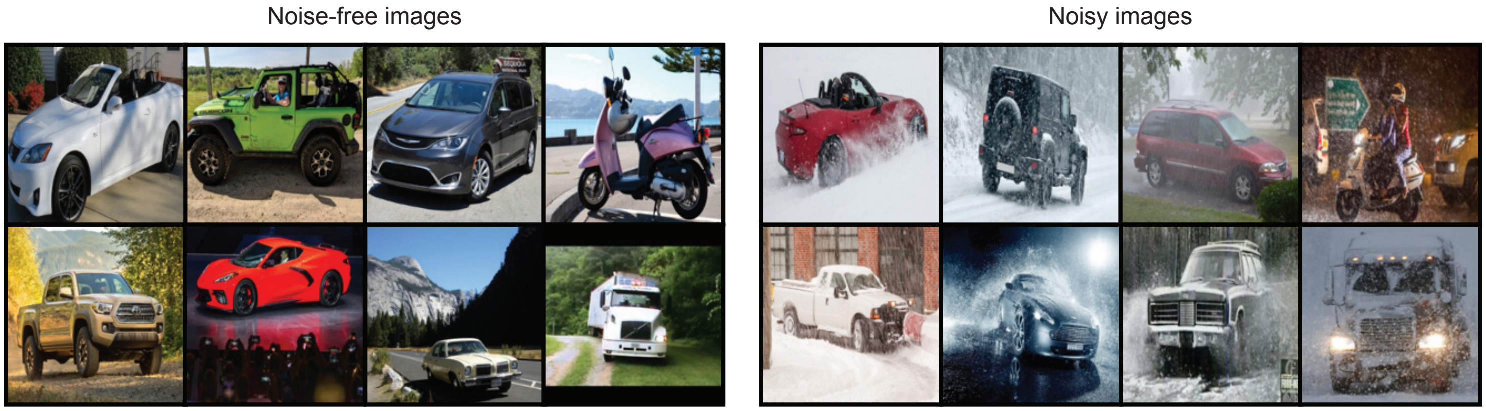


**Supplementary Figure 8.** Examples of real-world images of vehicles in noise-free and noisy conditions. Convertible, jeep, minivan, motor scooter, pick-up truck, sports car, station wagon, and trailer truck, from top-left to bottom-right.
